# Species sorting along a subsidy gradient alters community stability

**DOI:** 10.1101/031476

**Authors:** Mario E. Mucarella, Stuart E. Jones, Jay T. Lennon

**Affiliations:** Department of Biology, Indiana University, Bloomington, IN 47405 USA; Department of Biological Sciences, University of Notre Dame, Notre Dame, IN 46556 USA

**Keywords:** Resource Subsidies, Community Stability, Responsiveness, Microorganisms, Terrestrial DOC

## Abstract

The movement of resources between terrestrial and aquatic habitats has strong effects on ecological processes in recipient ecosystems. Allochthonous inputs modify the quality and quantity of the available resource pool in ways that may alter the composition and stability of recipient communities. Inputs of terrestrial dissolved organic carbon (tDOC) into aquatic ecosystems represent a large influx of resources that has the potential to affect local communities, especially microorganisms. To evaluate the effects terrestrial inputs on aquatic microbial community composition and stability, we manipulated the supply rate of tDOC to a set of experimental ponds. Along the tDOC supply gradient, we measured changes in diversity and taxon-specific changes in abundance and activity. We then determined community stability by perturbing each pond using a pulse of inorganic nutrients and measuring changes in composition and activity (i.e., responsiveness) along the gradient. Terrestrial DOC supply significantly altered the composition of the active microbial community. The composition of the active bacterial community changed via decreases in richness and evenness as well as taxon-specific changes in abundance and activity indicating species sorting along the gradient. Likewise, the responsiveness of the active bacterial community decreased along the gradient, which led to a more stable active community. We did not, however, observe these changes in diversity and stability in the total community (i.e., active and inactive organisms), which suggests that tDOC supply modifies microbial community stability through functional not structural changes. Together, these results show that altered aquatic terrestrial linkages can have profound effects on the activity and stability of the base of the food web and thus can alter ecosystem functioning.

## INTRODUCTION

Resource subsidies refer to the cross-boundary movement of materials and energy from donor to recipient ecosystems. These allochthonous inputs affect various ecological processes including population dynamics, species interactions, and ecosystem functioning (Polis et al. 1997, Holt 2004). Subsidies can also alter the stability of recipient communities. Theory suggests that low to moderate levels of subsidies stabilize recipient communities by promoting trophic omnivory (Polis et al. 1997, Huxel and McCann 1998) and dampening oscillations between consumers and their resources (DeAngelis 1992, Huxel and McCann 1998). The precise effect of subsidies on recipient communities, however, likely depends on the timing (Takimoto 2002), quality (Bartels et al. 2012), and quantity (Cottingham and Narayan 2013). Likewise, subsidies can also affect stability by altering the composition of recipient communities by selecting for and against particular species (Holt 2004). If subsides alter communities via species sorting, then they will also alter the distribution of species-specific traits that may affect stability, including resource specialization (Huxel and McCann 1998, Faria and Costa 2010) and metabolic plasticity (Comte et al. 2013). Together, the effect of subsidies on community stability is likely influenced by both subsidy properties (e.g., quantity) and consumer traits (e.g., resource specialization).

A well-recognized and pervasive subsidy is the movement of dissolved organic carbon (DOC) from terrestrial to aquatic ecosystems (Polis et al. 1997, Tranvik et al. 2009). In most inland water bodies, terrestrial DOC (tDOC) inputs exceed internal (i.e., autochthonous) inputs by aquatic autotrophs by at least an order of magnitude (Tranvik et al. 2009, Wilkinson et al. 2013). Additionally, there is growing evidence that tDOC inputs are increasing in many regions around the world owing to global change (Monteith et al. 2007), and it has been hypothesized that such changes in tDOC inputs will affect the functioning and stability of aquatic ecosystems (Jones and Lennon 2015, Solomon et al. 2015). Heterotrophic microorganisms are the primary consumers of tDOC in aquatic ecosystems. Despite having recalcitrant properties, tDOC is used by bacteria for anabolic and catabolic processes (Kritzberg et al. 2004, Lapierre et al. 2013). As such, heterotrophic bacteria are expected to mediate the aquatic ecosystem response to tDOC subsidies via changes in community composition and metabolic activity (Jones et al. 2009, Comte and del Giorgio 2011), and it has been hypothesized that, due to these changes, subsidies may alter the stability of recipient aquatic ecosystems (Wetzel 1999).

If subsidies modify the composition of recipient communities through processes such as species sorting, then subsidies could also alter the function and stability of recipient communities. Microbial communities are comprised of species with traits that link ecosystem functioning and community stability. For example, the degree to which communities are dominated by either generalist or specialists has important consequences for carbon cycling (Mou et al. 2008) and may explain how microbial communities respond to perturbations (Allison and Martiny 2008, Shade et al. 2011). Another important trait is metabolic plasticity or the ability of taxa to change physiological functions or to transition across levels of metabolic activity (e.g., active to dormant). Microbial communities consist of taxa that differ in activity and their degree of metabolic plasticity (del Giorgio and Gasol 2008, Lennon and Jones 2011), and the ability of individual taxa to rapidly adjust metabolic activity has been shown to be important for controlling ecosystem functions such as carbon and nitrogen cycling (Aanderud et al. 2015). In addition, the ability of taxa to adjust metabolic functions (Comte et al. 2013) or to transition across levels of activity (Lennon and Jones 2011) have been hypothesized to buffer communities against perturbations. Therefore, processes such as species sorting will alter the distribution of taxa and traits within communities and thus affect the stability and functioning of ecosystems.

In this study, we quantified the effects of tDOC supply on the diversity, composition, and stability of aquatic microbial communities. We hypothesized that subsidies would change bacterial composition via shifts in either the abundance or activity of taxa which would be reflective of species sorting. We further hypothesized that resource-driven shifts in composition would determine how subsidized bacterial communities respond to perturbations. To test our hypotheses, we manipulated tDOC supply rate in eleven experimental ponds. First, we used microbial community sequencing to assess changes in diversity and trends in abundance and activity across experimental treatments. Then, we measured community stability by quantifying changes in community composition in each pond following an inorganic nutrient perturbation. Results from our study provide a framework for how aquatic microbial communities may respond to increased resource subsidies such as tDOC, and show that subsidy-mediated shifts in composition alter the stability of communities.

## MATERIALS AND METHODS

***Experimental Ponds*** *–* We manipulated the supply rate of terrestrial dissolved organic carbon (tDOC) in eleven experimental ponds at the Michigan State University’s W.K. Kellogg Biological Station (KBS) Experimental Pond Facility. Each experimental pond has a 30 m diameter, a 2 m maximum depth, and an operating volume of approximately 10^6^ L. We established a tDOC supply gradient by adding different amounts of a commercially available source of humic substances (Super Hume, Crop Master USA) to each pond using a 5 horsepower trash pump. This source of humic substances is comprised of 17% humic acid and 13% fulvic acid and is known to be used by diverse heterotrophic bacteria (Lennon et al. 2013). We maintained the tDOC supply gradient for 100 days (6 June 2009 to 14 September 2009) by adding Super Hume to each pond on a weekly basis assuming a loss rate of 4.7 – 12.2% d^−1^ (Lennon et al., 2013; Jones & Lennon, 2015).

***Perturbation and Sampling*** *–* Nutrient limitation is typical for inland water bodies including the KBS experimental ponds, and aquatic communities are sensitive to episodic nutrient inputs (Elser et al. 1990). As such, nutrient pulses are a common perturbation to aquatic ecosystems and are often used in experiments to test questions about stability (Nowlin et al. 2008). We used an inorganic nutrient pulse to test the stability of aquatic microbial communities along the tDOC supply gradient. We added 500 L of an inorganic nutrient solution (NH_4_NO_3_ and Na_2_HPO_4_) to each experimental pond on day 82 (27 Aug 2009) using a 5 horsepower trash pump. The inorganic nutrient pulse elevated inorganic nitrogen (N) and phosphorus (P) concentrations of each pond by 565 μg L^−1^ and 50 μg L^−1^, respectively, while maintaining the initial N:P molar ratio. Prior to and after the inorganic nutrient pulse, we sampled each pond three times per week to track changes in water chemistry. We collected water samples from the center of each pond using a 1 m depth-integrated sampler. We measured DOC by oxidation and nondispersive infrared (NDIR) detection using a Shimadzu TOC-V carbon analyzer on 0.7 μm (Whatman, GF/F) filtered water samples. We measured total nitrogen concentrations spectrophotometrically after persulfate digestion using the second-derivative method (APHA 1998). We measured soluble reactive phosphorus concentrations colorometrically using the ammonium molybdate method (Wetzel and Likens 2000). Further details about chemical analyses and the nutrient perturbation can be found elsewhere (Jones and Lennon 2015).

***Bacterial Community Characterization*** *–* We characterized aquatic bacterial composition along the tDOC supply gradient using 16S rRNA sequencing approximately a week prior to (day 74) and after (day 88) the inorganic nutrient pulse. We collected water samples from the center of each pond using a 1 m depth-integrated sampler. We retained microbial biomass on 47 mm 0.2 μm Supor PES membrane filters (Pall) and stored immediately at −80 °C. Because microorganisms exist at various levels of metabolic activity which have differential effects on the structure and function of the community (del Giorgio and Gasol 2008), we used two approaches to characterizing composition. One approach used DNA, a stable molecule, to characterize the microbial community based on all taxa, regardless of activity level. We refer to the DNA approach as the “total community”. The second approach used RNA, an ephemeral molecule reflecting metabolic growth and activity (Elser et al. 2003), to characterize the microbial community based on the organisms that contribute to ecosystem function (Jones and Lennon 2010, Aanderud et al. 2015). We refer to the RNA approach as the “active community”. We extracted nucleic acids (DNA and RNA) using the PowerSoil Total RNA Extraction Kit with DNA Elution Accessory Kit (MoBio, Carlsbad, CA). We treated RNA extracts with DNase (Invitrogen) to degrade DNA prior to cDNA synthesis via the SuperScript III First Strand Synthesis Kit (Invitrogen). Once DNA and cDNA samples were cleaned and quantified, we amplified the 16S rRNA gene (DNA) and transcript (cDNA) using barcoded primers (515F and 806R) designed to work with the Roche 454 sequencing platform (Fierer et al. 2008; see Supplement for PCR sequences and conditions). We sequenced 16S rRNA amplicons at the Research Technology Support Facility at Michigan State University using the GS FLX Titanium chemistry. We processed raw 16S rRNA sequences using the software package *mothur* (version 1.32.1, Schloss et al. 2009). To identify operational taxonomic units (OTUs), we binned sequences into phylotypes using the Ribosomal Database Project’s 16S rRNA database and taxonomy as the reference (Cole et al. 2009).

***Community Diversity*** *–* To determine the effects of tDOC supply on bacterial community diversity, we measured alpha diversity in each pond prior to the inorganic nutrient pulse. First, we estimated taxonomic richness using a resampling approach. We subsampled communities to obtain a standardized 2000 observations per site and summed the number of OTUs represented in each subsample. We then resampled 999 additional times and calculated an average richness estimate (± SEM) for each site (Muscarella et al. 2014). We used Good’s Coverage to confirm that subsampling to 2000 observations was sufficient to make conclusions based on our data set (Good, 1953). Second, we determined taxonomic evenness using Simpson’s Evenness (Smith and Wilson 1996). Evenness was calculated using the same resampling approach we used for richness. We calculated richness and evenness for the total (DNA) and active (RNA) microbial communities. For each metric, we used an indicator variable multiple regression model (Neter et al. 1996) to test how diversity changed in response to tDOC supply with respect to both the total (DNA) and active (RNA) microbial community. In our regression model, we used supply rate as the continuous variable and community type (total vs. active) as the categorical variable. All statistical calculations were performed in the R computing environment (v 3.1.3, R Core Development Team 2012).

***Community Composition and Species Sorting*** *–* To determine the effects of tDOC supply on community composition and test for evidence of species sorting, we determined how bacterial communities and individual bacterial taxa responded to the tDOC gradient. First, we used PERMANOVA to determine if the bacterial community responded to tDOC supply rate for both the total and active communities (Anderson 2001). For each, if the PERMANOVA was significant we tested for evidence of species sorting defined here as species-specific directional changes in abundance (Jablonski 2008). We used Spearman’s rank-order correlation to measure the response of each taxon to tDOC supply. We identified responsive taxa based on significant rho-values after a Benjamini-Hochberg correction for multiple comparisons (Benjamini and Hochberg 1995). Positive rho-values indicate taxa that responded positively (in either the total or active community) to tDOC supply and negative rho-values identified taxa that responded negatively to tDOC supply. We then used the Ribosomal Database Project’s Taxonomy (Cole et al. 2009) and a curated freshwater bacteria database (Newton et al. 2011) to taxonomically identify each responsive taxa. All statistical calculations were performed in the R computing environment and PERMANOVA was implemented using the *adonis* function in the R package vegan (v 2.2–1; Oksanen et al. 2013). Taxonomic identifications were performed using *mothur.*

*Community Stability –* We determined the effects of the subsidy supply on community stability by calculating the change in community composition before and after the inorganic nutrient pulse (He et al. 1994, Brown 2003). First, we used Principal Coordinates Analysis (PCoA) to visualize differences in community composition based on Bray-Curtis distances. PCoA is a metric multidimensional scaling technique that allows objects to be oriented in a common reduced-dimension space while maintaining distance between objects as best as possible (Legendre and Legendre 2012). We used log_10_-transformed relative abundances and Bray-Curtis distance to reduce bias against low abundance taxa (Anderson et al. 2006, Legendre and Legendre 2012). We then measured the Euclidean distance in multivariate space between paired communities before and after the inorganic nutrient pulse using the first three multivariate axes (Brown, 2003). The Euclidean distance estimates the overall change in the microbial community, and we used the distance, which we refer to as “responsiveness”, as a metric of community stability (Grimm and Wissel 1997). Thus, a more stable community would be one that is less responsive to the inorganic nutrient pulse. We used an indicator variable multiple regression to test how subsidy supply rate altered community stability (i.e., responsiveness) with respect to both the total (DNA) and active (RNA) microbial communities. In our regression model, we used supply rate as the continuous variable and community type (total vs. active) as the categorical variable. All statistical calculations were performed in the R computing environment and PCoA was implemented using the *cmdscale* function in the R package vegan.

## RESULTS

*DOC Manipulation and Nutrient Perturbation* – The tDOC supply rates imposed (0 – 200 g C/m^2^) reflect the range of tDOC received by temperate lakes under current and future expected supply rates (Solomon et al. 2015), and established a DOC concentration gradient across ponds from 6 to 24 mg C/L (Supplemental Fig. 1). After the tDOC supply gradient had been established for 80 days, the inorganic nutrient pulse rapidly elevated nutrient concentrations approximately 10-fold while maintaining N:P ratios observed prior to the perturbation (Supplemental Fig. 2).

***Community Diversity*** *–* The tDOC supply gradient significantly decreased the richness and evenness of the active microbial community but had no effect on the diversity of the total community. An indicator variable multiple regression revealed that tDOC supply rate explained 85% of the observed variation in bacterial richness (F_3_, _15_ = 34.2, *P* < 0.001). Based on the total community (i.e., all taxa), richness did not change in response to tDOC supply (*P* = 0.50); however, the richness of the active community decreased in response to tDOC supply. Overall, we observed a 30% drop in the number of active taxa across the gradient (Fig. 1A; R^2^ = 0.64, *P* = 0.001). Similarly, an indicator variables multiple regression model revealed that tDOC supply rate explained 54% of the observed variation in community evenness (F_3_, _15_ = 8.12, *P* = 0.002). Based on the total community, community evenness did not change in response to tDOC supply (*P* = 0.32); however, the evenness of the active community decreased in response to tDOC supply with a 25% drop in evenness across the entire gradient (Fig. 1B; R^2^ = 0.51, *P* = 0.04).

**Fig. 1:**
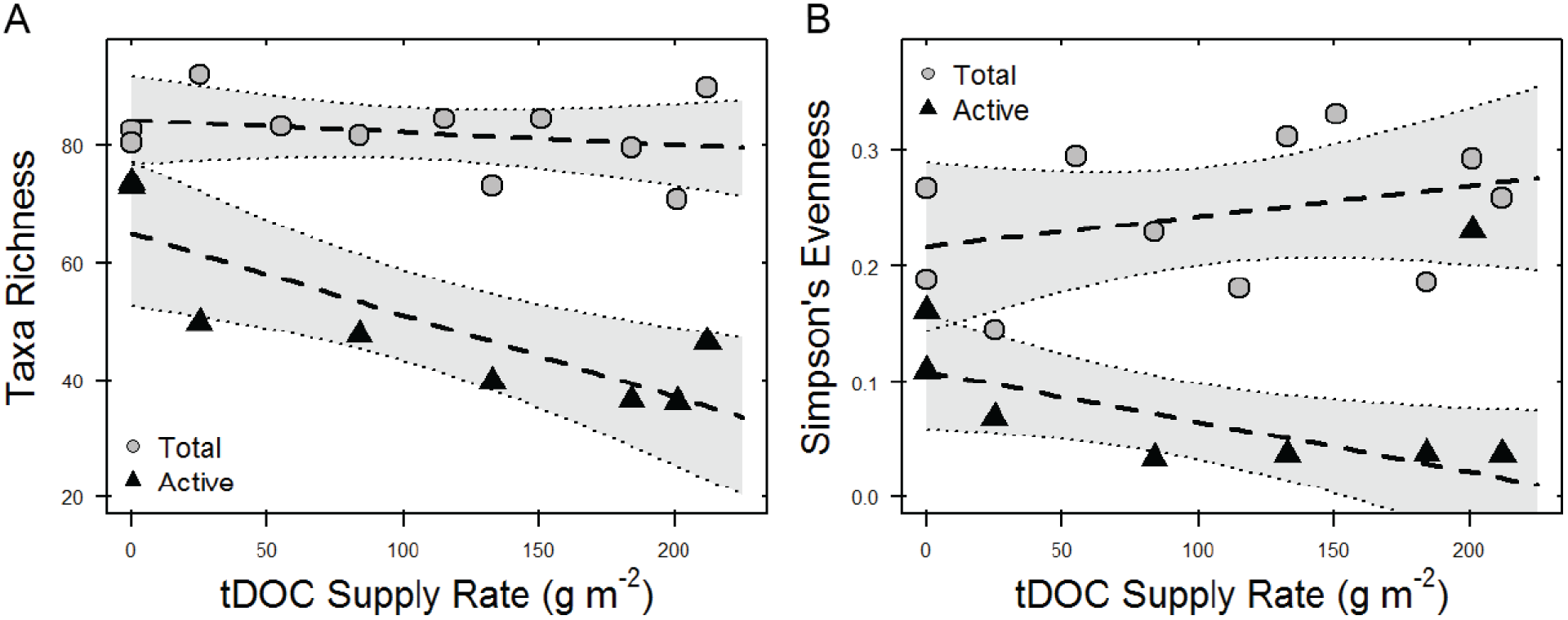
Community diversity estimates, richness (A) and evenness (B), for microbial communities in response to tDOC supply rate. For both, the diversity of the total community (DNA, circles) did not change in response to tDOC supply rate (A: *P* = 0.50; B: *P* = 0.32). However, both the richness and evenness of the active community (RNA, triangles) decreased in response to tDOC supply rate (A: *P* = 0.001; B: *P* = 0.04).

***Community Composition and Species Sorting*** *–* The tDOC supply gradient had a significant effect on community composition for both the total and active communities. Our PERMANOVA results show that tDOC supply altered the composition of the total (R^2^ = 0.17; *P* = 0.03) and active (R^2^ = 0.15, *P* = 0.04) bacterial community. In addition, we found evidence of species-specific responses in both the total and active communities. Together, 29 bacterial taxa (24% of the observed OTUs representing 26% of the total sequences) demonstrated a significant directional response based on the total (i.e., DNA) or active (i.e., RNA) community. Based on DNA analysis, 19 taxa significantly correlated with tDOC supply with eight positive and 11 negative relationships (Fig. 2B). Based on RNA analysis, 15 taxa significantly correlated with tDOC supply with one positive relationship and 14 negative relationships (Fig. 2C). The other 121 bacterial taxa did not demonstrate a significant relationship because their relative abundances were either constant or changed sporadically across the tDOC supply gradient.

**Fig. 2:**
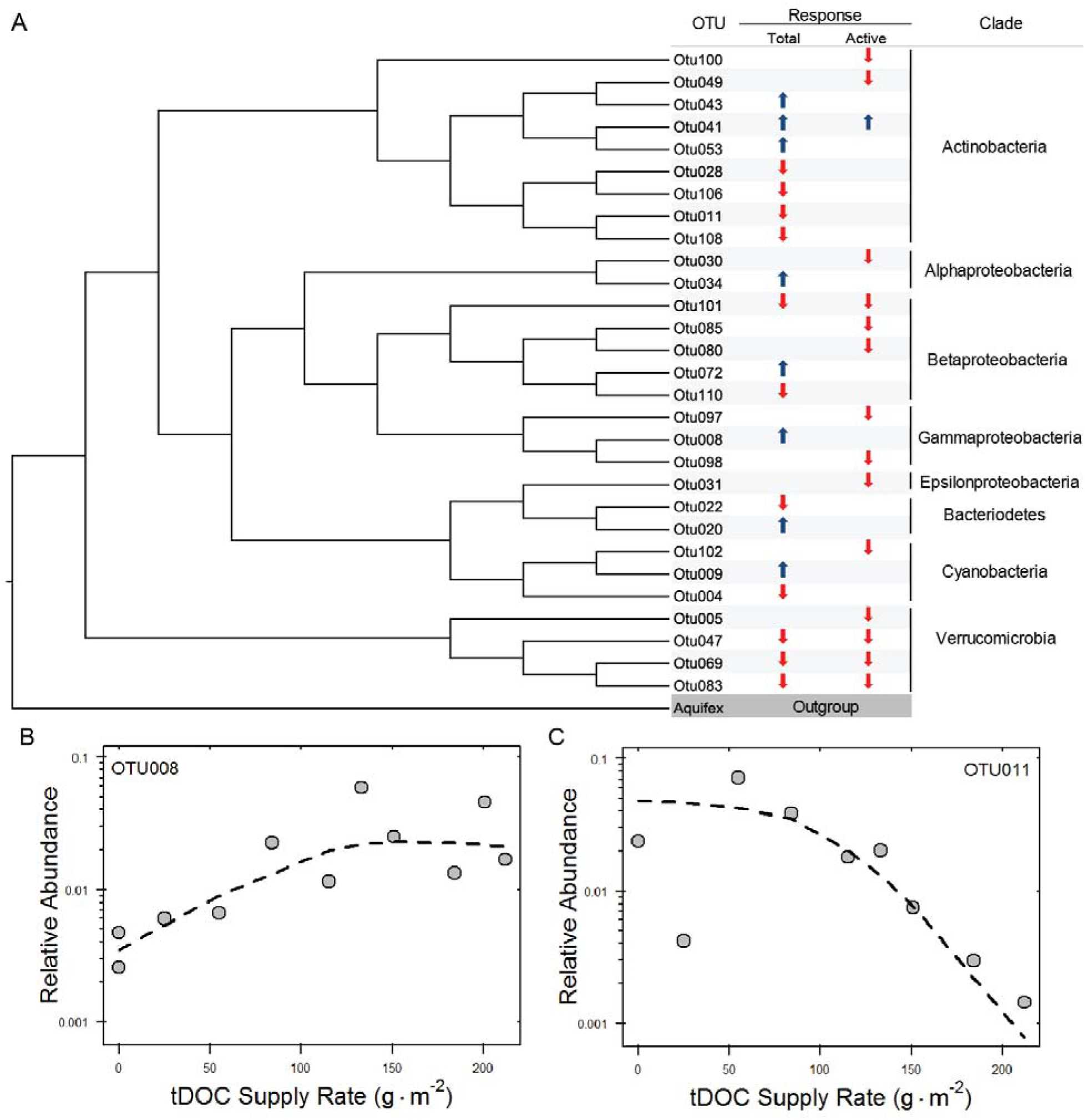
Taxon-specific changes in community composition along the tDOC supply rate gradient. A) Phylogenetic tree of responsive taxa as indicated by significant Spearman’s rank correlations. The consensus tree was inferred by using the Maximum Likelihood methods based on the Jukes-Cantor model, and 100 bootstraps using MEGA-6 (Tamura et al. 2013). Responses indicate significant increases (↑) or decreases (↓) in relative abundance in as tDOC supply rate increased. Responses are shown in regards to both the active and total communities. Clade refers to the phylogenetic group (phylum or subphylum) as inferred from best matches to the RDP taxonomy (Wang et al. 2007). Classifications based on known freshwater bacteria are found in Supplemental Table I. B) Example of significant positive correlation as shown by OTU008 *(Methylomonas* sp.). C) Example of a significant negative correlation as shown by OTU011 *(Rhodococcus* sp.). For each, log relative abundances are shown for OTU relative abundance in the total communities. Dashed lines indicate LOWESS regression trends and are included to demonstrate overall trends.

***Community Stability*** *–* The tDOC supply gradient significantly enhanced the stability of the active microbial community but had no effect on the stability of the total community. For our PCoA analysis, the first three ordination axes explained 60% of the variation between microbial communities across the tDOC supply gradient. An indicator variable multiple regression revealed that tDOC supply rate and community type (total or active) explained 83% of the observed variation in responsiveness (F_3_, _18_ = 35, *P* < 0.001). The responsiveness of the total community was low and did not change across the tDOC supply gradient (*P* = 0.37, Fig. 3A). However, the active community became less responsive as tDOC supply increased (*P* < 0.001, Fig. 3A), and there was a 35% drop in responsiveness across the entire gradient.

**Fig. 3:**
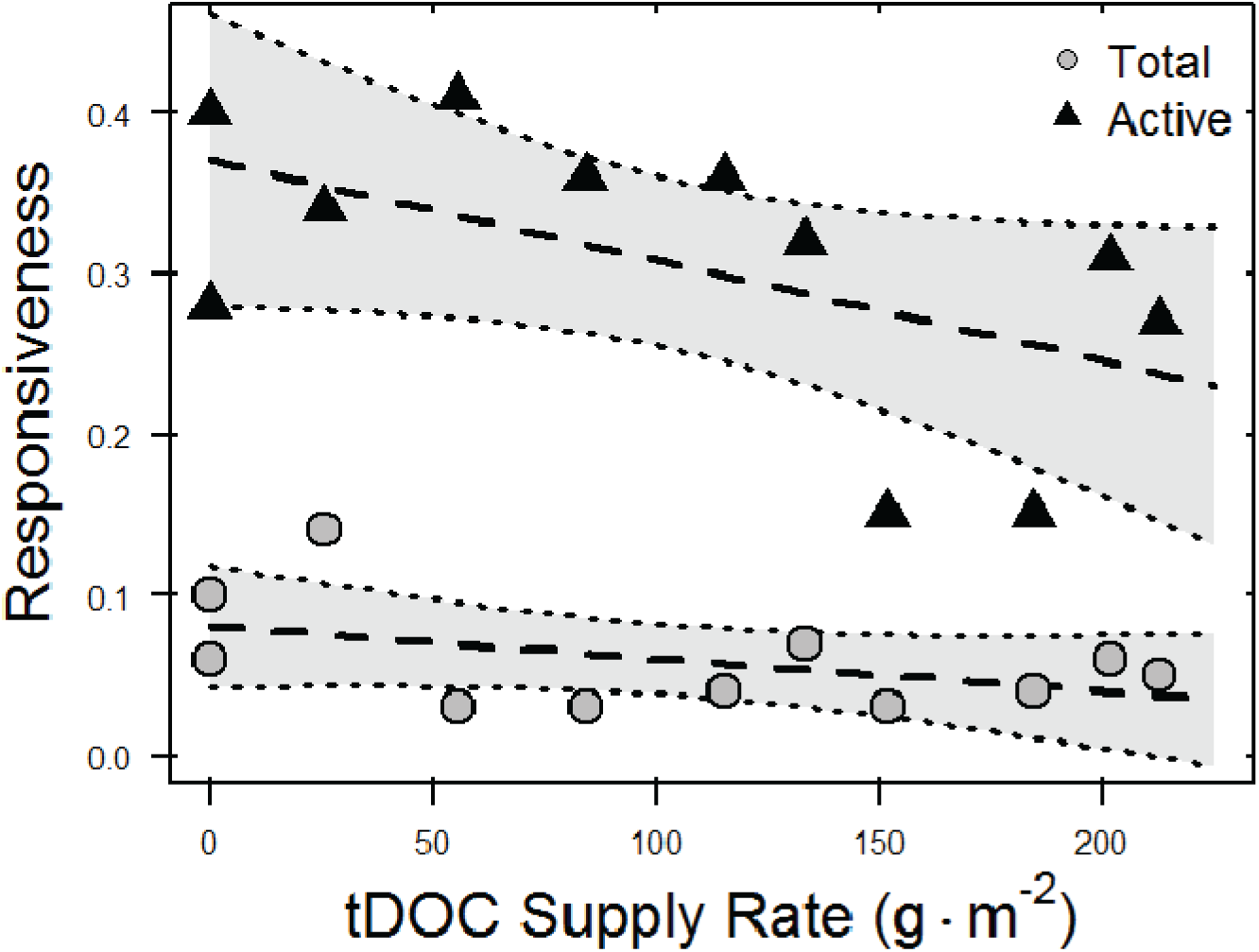
Community stability to an inorganic nutrient perturbation along the tDOC supply rate. Responsiveness was measured as the Euclidian distance between paired pre- and post-perturbation communities in multivariate space using the first three axes. Lower responsiveness indicates a more stable community. Based on the total community (DNA, circles),responsiveness did not change in as tDOC supply rate increased (*P* = 0.37). However, based on the active community (RNA, triangles), responsiveness decreased as tDOC supply rate increased (*P* < 0.001).

## DISCUSSION

The cross boundary movement of resources can alter the composition and stability of recipient communities, and therefore change community dynamics and ecosystem functioning. In this study, we manipulated the supply rate of terrestrial dissolved organic carbon (tDOC) to aquatic ecosystems and documented changes in the composition and stability of aquatic microbial communities. We found that the subsidy selected for and against certain taxa and suggest that this is evidence of tDOC-mediated species sorting. In addition, we found that supply rate increased the stability of the active members of the microbial community, and propose that via species sorting, tDOC established an active community that was less responsive to an inorganic nutrient pulse. Together, our results suggest that in the face nutrient perturbations, some subsidies (e.g. tDOC) select for specialized taxa that increase community stability. Furthermore, because these subsidies stabilize the organisms that regulate ecosystem functioning, the ability of ecosystems to contend with nutrient perturbations may diminish due to a reduced capacity for structural and functional responses by the members of the microbial community.

### Effects of terrestrial DOC on aquatic bacterial diversity

The tDOC supply gradient altered the aquatic microbial community by decreasing the richness and evenness of the active taxa. This suggests that ecosystems receiving high inputs of tDOC contain fewer active taxa and that certain taxa are disproportionately responsible for the majority of microbial activity in these ecosystems. There are multiple mechanisms by which increased tDOC inputs could decrease the diversity of active bacteria. First, tDOC is chromophoric and can decreases light availability, limit aquatic primary productivity, and thus reduces the concentration of labile, algal-derived resources that are used by many aquatic bacteria (Karlsson et al. 2009, Lennon et al. 2013, Jones and Lennon 2015). The decreased inputs of autochthonous inputs would have negative effects on aquatic microorganisms that rely on algal-derived resources (Kritzberg et al. 2004), and thus decrease the richness of the active community. Second, tDOC is a chemically and physically recalcitrant resource pool (Fellman et al. 2009), and it has been argued that only specialized consumers use tDOC (Fuchs et al. 2011). For example, the degradation of aromatic compounds requires specific metabolic pathways, most notably the beta-ketoadipate pathway (Fuchs et al. 2011). Therefore, tDOC subsidies may only favor taxa with specialized metabolic pathways (Mou et al. 2008, Fuchs et al. 2011). If these pathways are rare in the community then only few taxa would benefit from the chemical niches offered by tDOC. In sum, the disproportional benefit of tDOC would decrease the richness and evenness of the active microbial community.

### Species sorting alters community structure

In addition to changes in diversity, the altered resource environment modified the composition of aquatic bacterial communities by selecting for and against certain taxa (i.e. species sorting). We observed significant increases and decreases in taxa along the resource gradient. Although comparative studies have observed that DOC is an important factor structuring bacterial communities (Judd et al. 2006, Jones et al. 2009), here we present some of the first experimental evidence that tDOC supply can structure aquatic bacterial communities by species sorting.

In this study, we observed that some taxa increase in abundance in response to tDOC supply while other taxa decrease in abundance. For example, there was a 10-fold increase in the relative abundance of OTU008 (*Methylomonas* sp.) and a 10-fold decrease in the relative abundance of OTU011 (*Rhodococcus* sp.) across the gradient. There are multiple potential explanations for the taxon-specific changes in relative abundance we observed. First, the subsidy could be selecting against taxa unable to use tDOC while other taxa remain constant. For example, in addition to *Rhodococcus* (OTU011), four taxa belong to the Verrucomicrobia clade decreased in abundance and activity across the gradient (see Supplemental Table 1). In general, the Verrucomicrobia clade is thought to primarily use labile, algal-derived carbon (Newton et al. 2011). Second, taxa with beneficial traits (e.g., specialized metabolic functions) could be increasing in abundance while other taxa remain constant. For example, one of the strongest responding taxa in our study was an Actinobacterium (OTU043) related to the freshwater Actinobacteria tribe acSTL; in addition, we found increases in abundance for other Actinobacteria including members of the Luna3 and acIII tribes (see Supplemental Table 1). Though few cultured representatives exist, many members of the phylum Actinobacteria are known for the degradation of complex organic matter (Newton et al. 2011, Nelson and Carlson 2012). Last, because tDOC supply can promote additional microbial metabolisms, distinct groups of microorganisms may benefit from increased tDOC supply. For example, high concentrations of DOC may enhance lake methane cycling (Bastviken 2004, Lennon et al. 2006). We observed an increase in the relative abundance *Methylomonas* (OTU008) across the tDOC supply gradient, and this organism obtains carbon and energy from methane (i.e., methanotrophy). Together, our results provide evidence that subsidies, such as tDOC, structure bacterial communities via species sorting.

### Microbial Generalists and Specialists

In our study, we observed taxon-specific shifts in relative abundances, which indicate tDOC may favor taxa with specialized metabolic traits. Theory suggest that habitats with low concentrations of growth-limiting resources are thought to select for specialists that have lower minimal resource requirements, while habitats with higher concentrations of growth limiting resources may select for generalists (Wilson and Yoshimura 1994). In contrast, we increased the supply rate of tDOC, a low-quality complex resource requiring specialized metabolic traits, and observed taxon-specific increases in abundance. If these taxa are using tDOC as a resource, we can assume that they possess the required metabolic pathways and are specialists. Alternatively, these organisms may not be specialists. Work in marine environments has found that generalist microbes containing the diverse metabolic pathways are responsible for using tDOC (Mou et al. 2008, Newton et al. 2010). Likewise, work in the field of comparative genomics has shown that organisms that use large carbon resources tend to have larger genomes, which is a common signature of generalist life-history strategies among microorganisms (Livermore et al. 2014). It is possible, therefore, that some taxa contain specialized metabolic pathways but are in fact carbon substrate generalists rather than specialists. Therefore, we may need to change our expectations of the resource availability generalist-specialist framework to include resource complexity.

When resources are available to all consumers (low complexity) it is clear that resource availability will favor the consumers with the lowest minimal resource requirements (Tilman 1977); however, when resources are limited to consumers with unique traits (complex resources) then resource availability will favor consumers possessing these traits. Though often ignored, many nutrients – such as nitrogen, phosphorus, and carbon – exist as heterogeneous mixtures of molecules that differ in quality and bioavailability (Muscarella et al. 2014). As such, it is important to consider properties such as quality when predicting how resource subsidies will favor generalists versus specialists in a community.

### Community Response to Perturbations

In addition to compositional changes, tDOC supply rate altered how the bacterial community responded to an inorganic nutrient perturbation. There are four main predictions for how microbial communities will respond to perturbations: 1) resistant communities will show no changes; 2) resilient communities will quickly recover after perturbations and resume pre-disturbance structure and function; 3) sensitive, functionally redundant communities will change but maintain pre-disturbance function, and 4) sensitive, non-redundant communities will change in composition and function (Allison and Martiny 2008). Previous studies using nutrient perturbations have shown that microbial composition is generally sensitive to perturbation (Allison and Martiny 2008). This is not what we observed. In our study we found that the total bacterial communities had little to no response to a nutrient perturbation regardless of tDOC supply rate. This indicates that the total community was either resistant or resilient. However, we found that the active community changed across the tDOC supply gradient and that the active taxa responded less to the nutrient perturbation as tDOC supply increased. Because RNA is more ephemeral than DNA, we can assume that, if the community changed, the total community (based on DNA) would not return to pre-perturbation composition before the active community (based on RNA). Therefore, taking both the total and active communities into account, we argue that overall the total bacterial community was resistant to the inorganic nutrient pulse but the metabolically active taxa responded. In addition, the stability (i.e., resistance) of the metabolically active taxa increased as tDOC supply rate increased. We propose that the observed changes in the active community represent the functional responsiveness and metabolic plasticity of the bacterial community. Because only the functionally active organisms responded, the community response to the inorganic nutrient pulse was functional not structural; furthermore, because the functional responsiveness decreased as tDOC supply rate increased, subsidy inputs yielded a microbial community that was structurally and functionally stable.

### Subsidies and Community Stability

Overall, experimental tests to determine the effects of resource subsidies on community stability have been sparse (Nowlin et al. 2007). In this study, we found that a tDOC subsidy stabilized the recipient community by altering the composition of the microbial community through species sorting. One mechanism would be that the subsidy favored specialized taxa. However, due to their metabolic physiology, these specialized taxa may be relatively slow growing and unable to rapidly respond to nutrient pulses (Wetzel 1999). An alternative mechanism would be that the subsidy selected against fast-responding taxa. For example, at high tDOC supply shading limits primary productivity and the aquatic microbial community may receive less labile algal-derived carbon to promote fastidious, rapid responding taxa (Jones and Lennon 2015). Our data suggest that both of these mechanisms explain changes in composition due to tDOC supply. Therefore, we hypothesize that the quality of the subsidy influences community stability. Here, we added a low quality subsidy (tDOC) which selected for taxa with specialized metabolic functions and against fastidious taxa which would have been able to rapidly respond to the inorganic nutrient pulse. Conversely, systems receiving high-quality subsidies would favor more responsive taxa and thus communities would respond differently to perturbations. Therefore, subsidies affect community stability by altering the composition of the community through species sorting, but the community stability outcome will be dependent on the properties of the subsidy and the traits of the recipient consumers.

### Implications for Aquatic Ecosystems

Overall, our findings suggest that the active taxa within the microbial community are responsible for mediating the changes in composition and stability due to altered subsidy inputs. Because the active taxa control nutrient cycling, we also expect these changes to affect ecosystem function and stability. First, subsidy-induced changes in community composition will alter ecosystem functions. For example, the degree to which bacterial communities are physiologically flexible may have important consequences for ecosystem functions such as secondary productivity (Godwin and Cotner 2015), and our data suggest that tDOC subsidies may select for communities comprised of taxa that are less physiological flexibility and thus unable to rapidly respond to nutrient pulses. Second, changes in community stability may alter ecosystem stability. For example, enhanced community stability may yield reduced ecosystem stability because either functional or compositional changes are required for the ecosystems to respond to the nutrient perturbations (Comte and del Giorgio 2011). If inorganic nutrients from a pulse perturbation remain in the recipient ecosystem, due to slow rates of biological processing, ecosystem functioning will become destabilized (Cottingham and Carpenter 1994). In fact, previous work demonstrated that tDOC inputs diminished nutrient turnover time and thus destabilized aquatic ecosystem functioning (Jones and Lennon 2015). We hypothesized that the reduced ecosystem stability was due to a lack of microbial functional or structural responses. Our results confirmed this hypothesis. Together, our results support the view that subsidy supply alters species sorting in ways that alter the stability of recipient communities and ecosystems.

## ACKNOWLEDGEMENTS

This is Kellogg Biological Station contribution number KBS #1906 and this work was supported by the National Science Foundation (DEB-0842441 and DEB-1442246). We thank B.K. Lehmkuhl for technical assistance and M. Fisk and F. Guillemette for critical feedback on an earlier version of this manuscript. All sequence data and metadata have been submitted to NCBI and are available at BioProject PRJNA301893. All code for sequence processing and statistical analyses is available at https://github.com/LennonLab/SubsidyMicrobialStability.

